# Cerebellar contribution to cognitive deficits and prefrontal cortex dysfunction in Spinocerebellar Ataxia Type 1 (SCA1)

**DOI:** 10.1101/2024.07.10.602931

**Authors:** Kaelin Sbrocco, Ella Borgenheimer, Ying Zhang, Michael Koob, Marija Cvetanovic

**Affiliations:** Department of Neuroscience; Current affiliation: Baylor College of Medicine; Department of Lab Medicine and Pathology; Institute for Translational Neuroscience, University of Minnesota, 2101 6th Street SE, Minneapolis, MN 55455

## Abstract

The cerebellum’s role in cognition and its functional bi-directional connectivity with prefrontal cortex (PFC) is now recognized. However, how chronic cerebellar dysfunction affects PFC function and cognition remains less understood. Spinocerebellar ataxia type 1 (SCA1), is an inherited, fatal neurodegenerative disease caused by an abnormal expansion of glutamine (Q) encoding CAG repeats in the gene *Ataxin-1 (ATXN1*) and characterized by severe loss of Purkinje cells (PCs) in the cerebellum. Patients with SCA1 suffer from movement and balance deficits, cognitive decline and premature lethality. Cognitive deficits significantly impact patient’s quality of life, yet how exactly cerebellar degeneration contributes to cognitive deficits and PFC dysfunction in SCA1 is unknown. We have previously demonstrated that expression of mutant ATXN1 only in cerebellar Purkinje cells (PCs) is sufficient to cause cognitive deficits in a transgenic *ATXN1[82Q]* mouse line. To understand how cerebellar dysfunction impacts the PFC, we examined neuronal activity, synaptic density, and gene expression changes in the PFC of *ATXN1[82Q]* mice. Remarkably, we found decreased neuronal activity, reduced synaptic density, and altered expression of immediate early genes and pathways involved in glucose metabolism, inflammation and amphetamine in the PFC of *ATXN1[82Q]* mice. Furthermore, we characterized cellular and molecular PFC dysfunction in a novel conditional knock-in SCA1 line, *f-ATXN1^146Q^* mice, expressing floxed human expanded ATXN1 throughout the brain. Intriguingly, we found an increased number of neurons, increased synaptic density and large gene expression alterations in the PFC of *f-ATXN1^146Q^* mice. Finally, to precisely determine the role of cerebellar dysfunction in cognitive deficits and PFC dysfunction in SCA1, we crossed *f-ATXN1^146Q^* mice with *Pcp2-Cre* mice expressing Cre recombinase in PCs to delete *expanded ATXN1* only in PCs. Surprisingly, we have found that deleting *expanded ATXN1* in PCs exacerbated cognitive deficits and PFC dysfunction in these mice. Our findings demonstrate that circumscribed cerebellar dysfunction is sufficient to impact PFC activity and synaptic connectivity impairing cognition. However, when multiple brain regions are impacted in disease, cerebellar dysfunction may ameliorate PFC pathology and cognitive performance.

## Introduction

The prefrontal cortex (PFC) occupies the anterior portion of the cerebral cortex and plays a crucial role in a wide range of cognitive functions and behaviors^1^. Higher order processes such as decision making, problem solving, working memory, strategy formation, and social cognition have been shown to involve the PFC^2,3^. Given its complex functions and extensive connectivity to other brain regions, the PFC is often implicated in the pathology of neurological and psychiatric disorders^4,5^.

Imaging, clinical and pre-clinical, anatomical and functional connectivity studies all provide strong evidence for the functional interaction of the PFC with the cerebellum and the role of the cerebellum in cognition^6–9^. How cerebellar dysfunction impacts the PFC and cognitive impairments in disease is less clear. Spinocerebellar ataxia type 1 (SCA1) is a progressive neurodegenerative disease caused by the abnormal expansion of glutamine (Q) encoding CAG repeats in the *Ataxin-1* (*ATXN1*) gene^10^. SCA1 belongs to a group of polyglutamine (polyQ) diseases which also include Huntington’s disease, spinobulbar muscular atrophy (SBARNES MAZEA), SCA2, 3, 6, 7, 17, and dentatorubral pallidoluysian atrophy (DRPLA)^11,12^. Patients with SCA1 suffer from impaired balance and coordination, as well as cognitive and affective deficits, most predominantly in the areas of executive function and depression^13–21^. Postmortem pathology and imaging studies revealed a profound loss of Purkinje cells (PCs) in the cerebellum, and less severe but wide spread degenerative changes in the brain, including the cerebral cortex ^22,23^. While the PFC has been implicated in cognitive and affective functions, including executive function, PFC pathology in SCA1 is little understood. Examining cellular and molecular pathology in the PFC is necessary not only to increase our understanding of the PFC pathogenesis and cognitive decline in SCA1, but also of how degeneration in the cerebellum affects the PFC.

Mouse models are very fruitful in increasing our understanding of the mechanisms of neurodegeneration and this is very true for SCA1 pathogenesis^12,24,25^. In previous studies, we demonstrated that expression of mutant ATXN1 only in PCs is sufficient to impair performance in Barnes Maze and fear conditioning assays in *ATXN1[82Q]* mice^26^. As the PFC plays an important role in strategy development, we first asked how cerebellar dysfunction in *ATXN1[82Q]* transgenic mice impacts the PFC on a cellular and molecular level. We then investigated PFC alterations in a novel SCA1 conditional knock-in line^27^, *f-ATXN1^146Q^* mice. Finally, we deleted expanded ATXN1 only in PCs to directly determine the impact of PC-specific expression of mutant ATXN1 on PFC pathology and cognitive deficits in SCA1.

## Materials and Methods

### Mice

Mice were housed in a temperature-and humidity-controlled room on a 12 hour light/12 hour dark cycle with access to food and water *ad libitum*. Mice were tested starting from 12 weeks of age. All animal experiments were performed in compliance with the National Institutes of Health’s Guide for the Care and Use of Laboratory Animals and approved by the University of Minnesota Institutional Animal Care and Use Committee In all experiments, equal numbers of male and female mice were used. All mice were age matched within experiments, and littermate controls were used when possible. The *ATXN1[82Q]* mice, *Pcp2-Cre,* and *f-ATXN^146Q^* mice were gifts from Dr. Harry Orr and Michael Koob. *ATXN1[82Q]* mice were created originally on FVB background and were backcrossed onto a C57/Bl6 background. In the *Pcp2-Cre* mice^28^, Cre expression is limited to PCs in the cerebellum, resulting in selective deletion of mutant ATXN1 in PCs in *f-ATXN1^146Q^*;*Pcp2-Cre* mice.

### Cognitive testing

Sample sizes in the behavioral tests were determined using power analysis and the prior experience with these tests, or previous reports using similar methodology. Experimenters were blinded to the genotype during all tests.

*Barnes Maze.* The maze was a white circular platform 91 cm in diameter with 20 5-cm circular holes spaced evenly around the edge, raised approximately 92 cm above the floor. One of the holes led to a 5 cm wide x 11 cm long x 5 cm deep opaque box (the “escape box”) and the other 19 were covered. The testing room had visual cues on the walls to serve as landmarks, and all objects in the room were kept in the same places for every trial. The position of each mouse was tracked using AnyMaze software. Mice were exposed to the maze for four 3-minute trials per day during four consecutive training days (intertrial interval of approximately 15 minutes). Mice which did not enter the escape box within 3 minutes were gently guided to it. Training day data is reported as a path length (distance traveled before entering the escape hole) and analyzed by two-way repeated measures ANOVA. A probe test was conducted 24 hours after the last training session. For the probe test, the escape hole was covered and each mouse was allowed to explore the maze freely for 90 seconds. The time spent in each quadrant of the maze was recorded, and the amount of time spent in the goal quadrant (the quadrant centered on the goal hole) was analyzed by one-way ANOVA.

Search strategies on the training days were automatically classified and assigned cognitive scores using the Barnes Maze unbiased strategy (BUNS) classification tool as described by Illouz *et al.*^22^ In order to compare learning rates between groups, cognitive scores for each mouse on each of the 16 training trials were plotted in GraphPad Prism 7.0. Linear regression was performed for each group and the slopes and elevations of the lines were compared using Prism’s Analysis function.

In one cohort of mice, wild-type mice did not demonstrate learning. As this indicates technical problems, we have excluded that whole cohort from the analysis.

*Contextual fear conditioning.* Conditioning took place in chambers with a floor consisting of stainless steel rods through which shocks were delivered (Med Associates #ENV-008-FPU-M). On day 1, mice were placed in the chambers for a 10-minute period during which they received five foot shocks (0.70 mA, 2-second duration). Freezing during the 60 seconds after each shock was quantified automatically using VideoFreeze software (freezing was defined as a motion index ≤15 lasting ≥500 ms). 24 hours after the initial conditioning, mice were returned to the same chambers with the shock generators turned off and freezing behavior was monitored for 3 minutes. 1-2 hours after being placed in the conditioned context, mice were placed in a second context for 3 minutes to measure baseline freezing. The baseline context used the same chambers but differed from the conditioned context in floor texture (smooth plastic versus metal rods), shape (curved plastic wall versus square metal wall), and odor (0.5% vanilla extract versus 33% Simple Green). Acquisition of freezing responses is reported as percent freezing in the 60-second period following each of the 5 foot shocks, analyzed by two-way repeated measures ANOVA. 24-hour recall is reported as percent freezing in each context over the 3-min test period, analyzed by two-way repeated measures ANOVA.

### Prefrontal cortex dissection

To dissect the PFC, isolated brains were placed into a coronal sectioning brain matrix with the dorsal side up. One razor blade was placed along the sagittal midline, making sure that the blade pushes all the way to the bottom of the matrix. An additional two razor blades were consequently positioned 2mm apart from each other in the left hemisphere coronally. One blade was placed between the olfactory bulbs and the start of cortical tissue. The other was positioned 2mm from that blade caudally. The most dorsal medial quadrant of this isolation was then separated and claimed as PFC.

### RNAseq

RNA was extracted from dissected regions using TRIzol Reagent (Life Technologies). RNA was sent to the University of Minnesota Genomics Center for quality control, including fluorometric quantification (RiboGreen assay, Life Technologies) and RNA integrity with capillary electrophoresis (Agilent BioAnalyzer 2100, Agilent Technologies Inc.). All samples with RNA integrity numbers (RINs) 8.0 or greater proceeded to RNA sequencing on an Illumina NextSeq 550 using a 100-nt paired-end read strategy^29^. Data is stored on University of Minnesota Supercomputing Institute Servers. To analyze data raw, paired short reads were analyzed through CHURP (Collection of Hierarchical UMII- RIS Pipelines, PEARC’19 Proceeding, Article No. 96), which includes data quality control via FastQC, data preprocessing via Trimmomatic (Bolger et al., 2014), mapping via HiSat2 against reference mouse genome and expression quantification via Subread. Resulting count matrix of gene expression was used as input of R (https://www.r-project.org/) package: EdgeR (v3.32.1), which tested the differential gene expression and visualized the behavior of expected invariant control genes and published positive control genes. Genes with resulting FDR values less than 0.05 were considered significant. EdgeR was used to reveal sample outliers through Multi-Dimensional Scaling (MDS) and Principle Component Analysis (PCA). Pathway analysis and dot plot creation was preformed using the clusterProfiler package (v3.18.1) using the top 500 DEGs based on absolute value of LogFC. Heatmaps were created using pheatmap package (https://CRAN.R-project.org/package=pheatmap)(v1.0.12). Volcano plots were created using BiocMananger (https://CRAN.R-project.org/package=BiocManager) (v1.30.16) and EnhancedVolcano (https://github.com/kevinblighe/EnhancedVolcano) (v1.8.0) packages. Pathway activity score was calculated using R package GSVA, we then averaged the score across samples from the same brain region with the same genotype.

### Immunofluorescence

IF was performed on a minimum of six different floating 40-μm-thick brain slices from each mouse (six technical replicates per mouse per region or antibody of interest). We used primary antibodies against c-Fos (rabbit, Abcam, ab190289), neuronal marker neuronal nuclei (NeuN) (rabbit, Abcam, Ab104225), PSD95 (rabbit, Thermo Fisher Scientific, 51-69000), vesicular glutamate transporter 2 (VGLUT2) (guinea pig, Millipore, AB2251-I), Gephyrin (rabbit, Thermo Fisher Scientific, PA5-29036), and VGAT (guinea pig, Synaptic Systems,131005) as previously described^29,30^.

### Imaging and image analysis

For NeuN and cFos image acquisition, confocal images were acquired using a confocal microscope (Olympus FV1000) using a 20X oil objective. Z-stacks consisting of twenty non-overlapping 1-μm-thick slices were taken of each stained brain slice per PFC (i.e., six z-stacks per mouse, each taken from a different brain slice). To quantify total neuron count, we counted the number of NeuN+ puncta using the ImageJ Analyze Particles feature. To quantify relative neuronal activity, we counted the number of cFos+ puncta using the ImageJ Analyze Particles feature. Data is represented as the density of NeuN+ or cFos+ respective to the PFC area analyzed. For synaptic quantification image acquisition, confocal images were acquired using a confocal microscope (Leica Stellaris 8) using a 63X oil objective. Three Z-stacks consisting of fifteen non-overlapping 0.34-μm-thick slices were taken of each stained brain slice per PFC (i.e., eighteen z-stacks per mouse, three z-stacks per brain slice). To quantify excitatory and inhibitory synaptic content, we analyzed synaptic quantification images using the ImageJ Puncta Analyzer plugin. Presynaptic (vGLUT2 and VGAT) signal and postsynaptic (psd95 and Gephyrin) signal were quantified individually. Colocalized puncta are then counted based on the Puncta Analyzer parameters.

### Statistical Analysis

Statistical tests were performed using GraphPad Prism. Data was tested for normal distribution using Kolmogrov-Smirnov and Shapiro-Wilk tests. Parametric tests were performed if normal distribution of the data was established, otherwise non-parametric tests were chosen. Data was analyzed using two-way ANOVA, one-way ANOVA followed by either Tukey’s HSD or Bonferroni post-hoc test, or Student’s t-test.

## Results

### Cerebellar expression of mutant ATXN1 is sufficient to impact neuronal activity, synaptic connectivity, and gene expression in the PFC of ATXN1[82Q] mice

To understand how mutant ATXN1 (expanded ATXN1) expression in PCs impacts the PFC at a cellular level, we quantified neuronal density in the PFC of *ATXN1[82Q]* mice and littermate controls (Supplementary Figure 1). We did not find any change in the density of NeuN+ neurons in the PFC of 18 weeks old *ATXN1[82Q]* mice (Supplementary Figure 2A). This was not surprising as there is minimal loss of PCs in the cerebellum of these mice at 18 weeks^31,32^. However, even at 28 weeks when there is a detectable loss of PCs in *ATXN1[82Q]* mice, the density of PFC neurons was not altered (Figure 1A).

**Figure 1.**
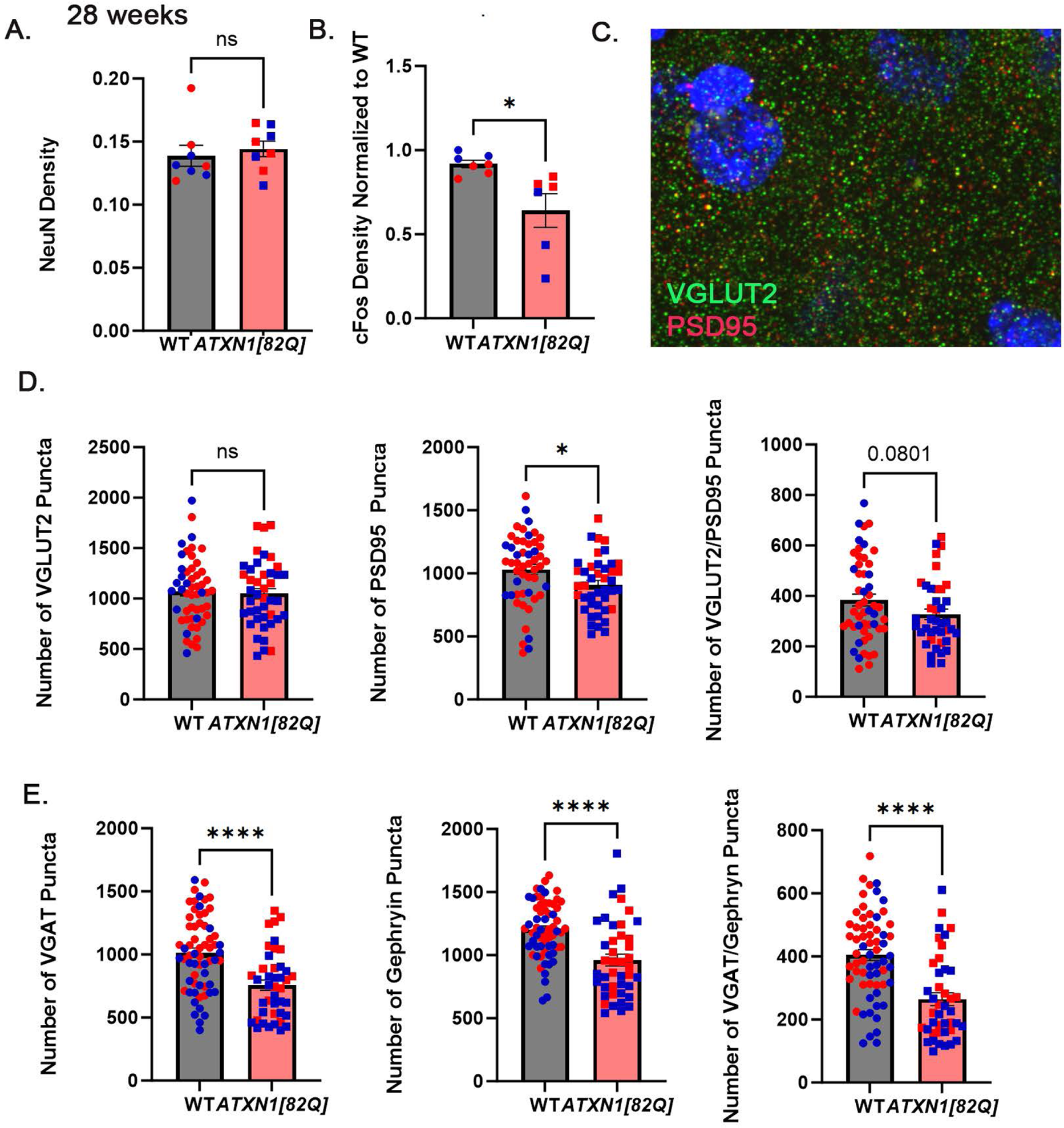
PFC pathology in SCA1 transgenic mice. Saggital brain sections of 26 week old *ATXN1[82]* mice and their wild-type littermates were immunostained for NeuN, c-Fos, VGLUT2, PSD95, VGAT and gephyrin, and confocal z-stack images were obtained. A. Quantification of NeuN density. B. Quantification of c-Fos density. C. Image of VGLUT2 and PSD95 puncta. D. Quantification of excitatory presynaptic VGLUT2+, postsynaptic PSD95+ and co-localized VGLUT2/PSD95 puncta. E. Quantification of inhibitory presynaptic VGAT+, postsynaptic Gephyrin+ and co-localized VGAT/gephyrin puncta. N= 6-12 *ATXN1[82]* mice and their wild-type littermates. * P< 0.05, **** P<0.0001 Student’s t-test.

Next, we quantified neuronal activity in the PFC using a molecular marker, c-Fos^33^. By counting the number of c-Fos-positive cells in the PFC, we found a significant effect on c-Fos levels in the PFC (Figure 1B). Mutant ATXN1 expression in cerebellar Purkinje cells resulted in a decreased c-Fos expression in the PFC of *ATXN1[82Q]* mice compared to wild-type littermate controls. We next asked whether this decrease in cFos expression *ATXN1[82Q]* mice experienced was caused by altered synaptic inputs. Using pre-and post-synaptic markers of excitatory (VGLUT2 and PSD95) and inhibitory (VGAT and gephyrin) synapses we quantified the number of labeled puncta and their co-localization in the PFC of *ATXN1[82Q]* mice and wild-type littermate controls(Figure 1C)^30^. We have found a significant decrease in the excitatory post-synaptic PSD95 puncta and a trending decrease in the number of VGLUT2/PSD95 co-localized puncta (Figures 1 D).

Density of VGAT and gephyrin puncta, as well as VGAT/Gephyrin co-localization were significantly decreased in the PFC of *ATXN1[82Q]* mice (Figures 1W).

To understand how expanded ATXN1 expression in PCs impacts the PFC at a molecular level, we next investigated genes altered in the PFC of *ATXN1[82Q]* mice. We dissected the PFC from *ATXN1[82Q]* mice and wild-type littermates, isolated RNA, and performed RNA sequencing. We found 341 differentially expressed genes (DEGs, defined as absolute log2FC > 0.6 and p < 0.05) in the PFC of *ATXN1[82Q]* mice, with majority of genes (∼ 73%, 251 DEGs) upregulated (Figure 2A, Table 1). We found increased expression of genes *Slc2a2*, and *Slc2a4* encoding for neuronal glucose transporters GLUT2 and GLUT4. GLUT2 has low affinity for glucose, is considered a glucose sensor in the brain, and is involved in control of sugar consumption^34^. GLUT4 is an insulin-responsive glucose transporter thought to mediate the effects of insulin or insulin-like growth factor on cognition^35^. Expression of transcriptional factors was also impacted, including decreased expression of immediate early genes *Arc*, *Fosb*, and *Jun*. We also found increased expression of *Eomes*^3^, developmental cortical neurogenesis regulator*, Egr4*^37^, a regulator of the neuronal differentiation and maintenance state, and *Egr2,* known as regulator of *FosB* and *Jun*^38^. Intriguingly, expression of several neuroinflammatory genes including *Ccl17, Ccl28, Il3a, Nlrp3, Nlrp12, Nfkbid* and *Il1bos* was also altered in the PFC of ATXN1[82Q] mice.

**Figure 2.**
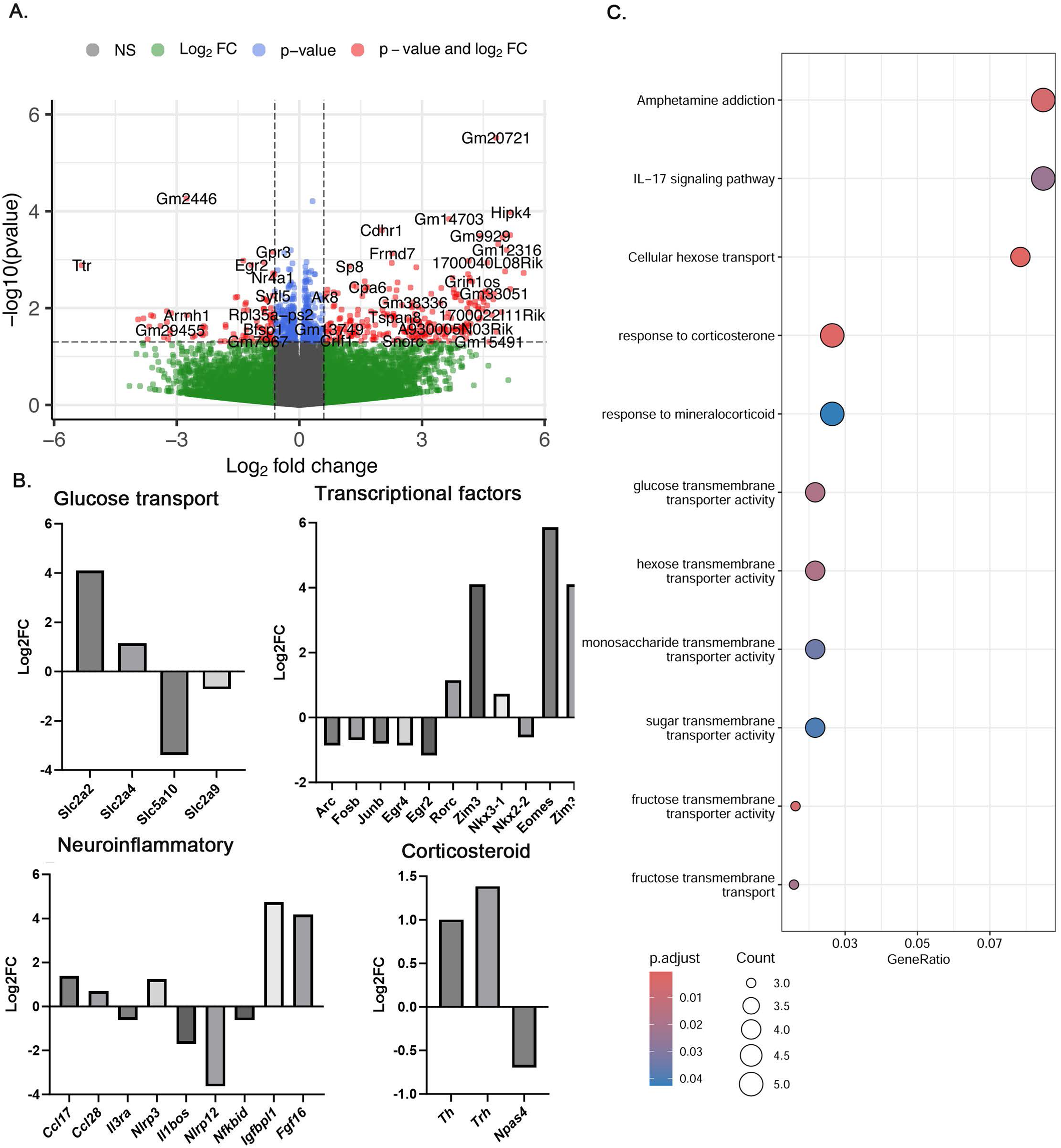
Gene expression changes in PFC of *ATXN1[82Q]* mice. A. Volcano Plot of Differentially Expressed genes (DEGs) between WT and *ATXN1[82Q]* PFC ( N= 6 each). B. Altered expressio of genes involved in glucose transport, transcriptional factors, neuroinflammation, and corticosterol signaling. For all genes p < 0.05. C. Pathway analysis indicating altered signaling in PFC of *ATXN1[82Q]* mice.

Gene ontology (GO) pathway analysis identified alterations in monosaccharide transporter activity and response to corticosterone. Kyoto Encyclopedia of Genes and Genomes (KEGG) analysis identified alterations in inflammatory IL17 pathway and amphetamine addiction (Figure 2C). Among the corticosterone pathways genes were *Th*^39^, *Trh*^40^ and *Npas*^41–44^.

These results suggest that mutant ATXN1 expression in PCs of the cerebellum is sufficient to cause adaptations in the PFC including altered gene expression, reduced synaptic density and neuronal activity in the PFC.

### PFC dysfunction resulting from the ubiquitous expression of expanded ATXN1 in SCA1 knock-in mice includes changes in PFC neurons, neuronal activity, synaptic connectivity, and transcriptomic dysregulation

While there is a profound loss of PCs in the cerebellum of patients with SCA1, less severe pathology has been reported in other brain regions, including the cortex^13,22^. ATXN1 is expressed in different cell types throughout brain (https://www.proteinatlas.org/ENSG00000124788-ATXN1/brain), and our RNAseq data^45^ has shown robust expression of *Atxn1* in the prefrontal cortex (Supplementary Figure 3). We next investigated how expression of mutant ATXN1 beyond PCs contributes to the PFC pathology in a SCA1 knock-in line, *f-ATXN1^146Q^* mice, that express mutant ATXN1 throughout the brain under an endogenous promoter^27^.

Intriguingly, we found a significant increase in neuronal density in the PFC of *f-ATXN1^146Q^* mice at 18 weeks of age (Figure 3A). Quantification of neuronal activity revealed a trending decrease in c-Fos expression in the PFC of 18-week-old *f-ATXN1^146Q^* mice (Figure 3B-C). We found a significant increase in the density of pre-and post-synaptic excitatory VGLUT2 and PSD95 puncta, as well as, an increased VGLUT2/PSD95 colocalization indicative of increased density of excitatory synapses in the PFC of *f-ATXN1^146Q^* mice (Figure 3D-F). Density of inhibitory presynaptic VGAT puncta was also increased in the PFC of *f-ATXN1^146Q^* mice (Figure 3G).

**Figure 3.**
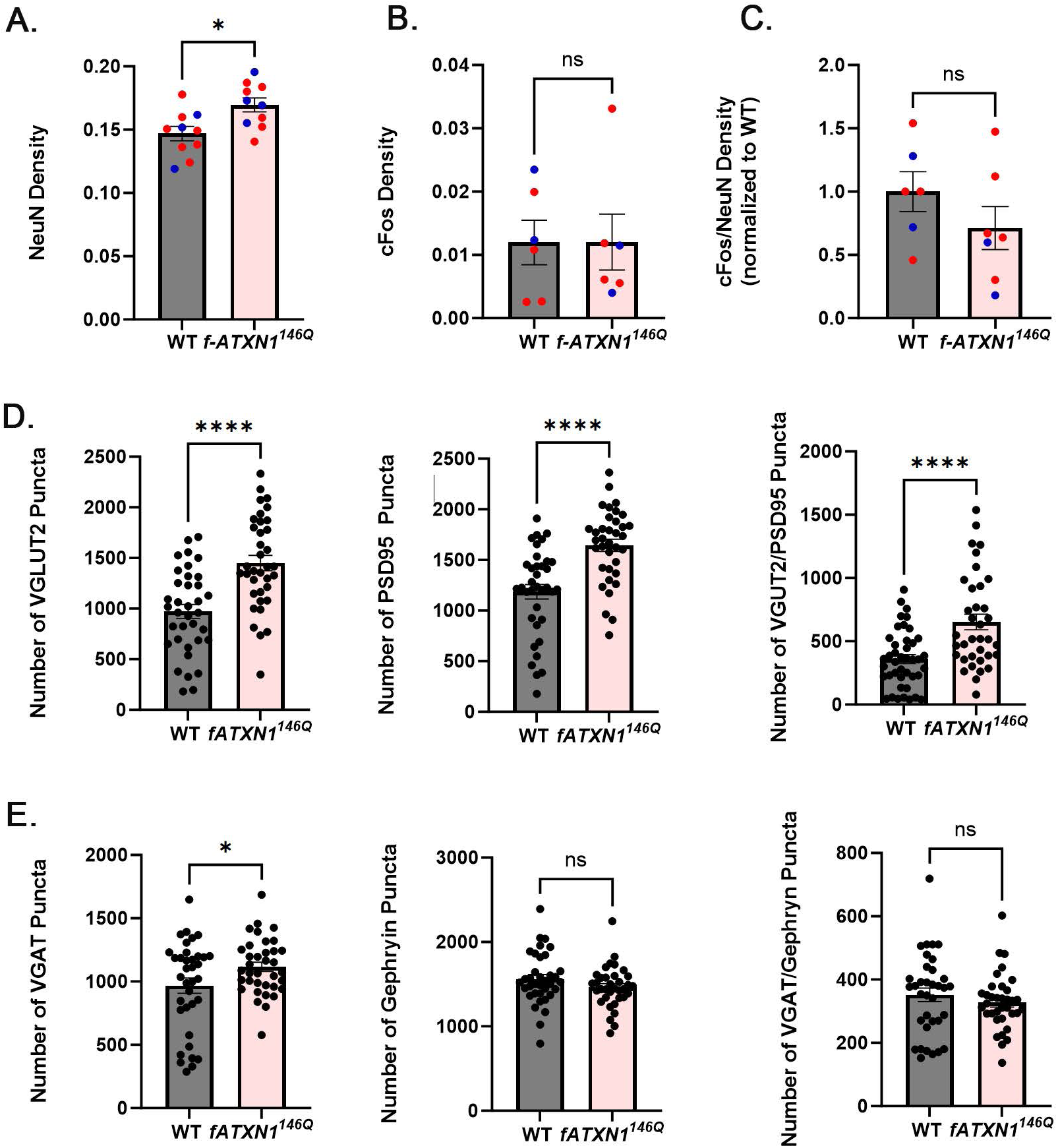
PFC pathology in *f-ATXN1^146Q^* mice. Saggital brain sections of 18-week-old *f-ATXN1^146Q^* mice and their wild-type littermates were immunostained for NeuN, c-Fos, VGLUT2, PSD95, VGAT and gephyrin, and confocal z-stack images were obtained. A. Quantification of NeuN density. B. Quantification of c-Fos density. C. Image of VGLUT2 and PSD95 puncta. D. Quantification of excitatory presynaptic VGLUT2+, postsynaptic PSD95+ and co-localized VGLUT2/PSD95 puncta. E. Quantification of inhibitory presynaptic VGAT+, postsynaptic Gephyrin+ and co-localized VGAT/gephyrin puncta. N= 6-12 *f-ATXN1^146Q^* mice and their wild-type littermates. * P< 0.05, **** P<0.0001 Student’s t-test.

To investigate how disease progression impacts these PFC adaptations in SCA1 knock-in mice, we quantified neuronal and synaptic density and neuronal activity in PFC of *f-ATXN1^146Q^* mice at a later disease stage (28 weeks). There was no difference in the number of NeuN+ neurons in the PFC of 28-week-old *f-ATXN1^146Q^* mice and their wild-type littermates (Supplementary Figure 3). We also found a significant effect on c-Fos levels in the PFC of 28 –week-old *f-ATXN1^146Q^* mice, with decreased c-Fos expression compared to wild-type controls, and a significant decrease in the density of inhibitory post-synaptic gephyrin puncta (Supplementary Figure 4).

To explore molecular changes in the PFC caused by ubiquitous mutant ATXN1 expression we performed RNAseq on the PFC dissected from *f-ATXN1^146Q^* mice. We have found a large transcriptional dysregulation with 1333 DEGs in the PFC of *f-ATXN1^146Q^* mice (Figure 4A, Table 2). Most of the genes (817 DEGs, or 61.2%) were downregulated, which is consistent with the known role of ATXN1 in gene repression^45,46^. Amongst the downregulated DEGs were many synaptic genes, such as *Snap25* encoding for protein SNAP25 which is part of the plasma membrane SNARE complex critical for neurotransmitter release as well as genes encoding for postsynaptic proteins including *Dlg4,* encoding for PSD95 and *Homer1* (Figure 4B). We found reduced expression of neurotransmitter receptors including encoding for GABAR (*Gabbr2* and *Gabrd*), glutamatergic (*Grm2*, *Grm3* and *Grm4*), cholinergic (*Chrna 2, 4, 5*, *Chrnb3, Chrm1, 3. 4*), dopamine (*Drd5*) and opioid receptors (*Oprd1* and *Oprk1*), while two genes encoding serotonin receptors *Hrta3* and *Htr5d* were slightly upregulated (Figure 4B). Expression of both excitatory and inhibitory neuron markers were also significantly decreased (Figure 4B). Many neuroinflammatory genes were also altered and when we examined expression of genes thought to represent M1 and M2 phenotypes we have found that most were downregulated (Figure 4B).

**Figure 4.**
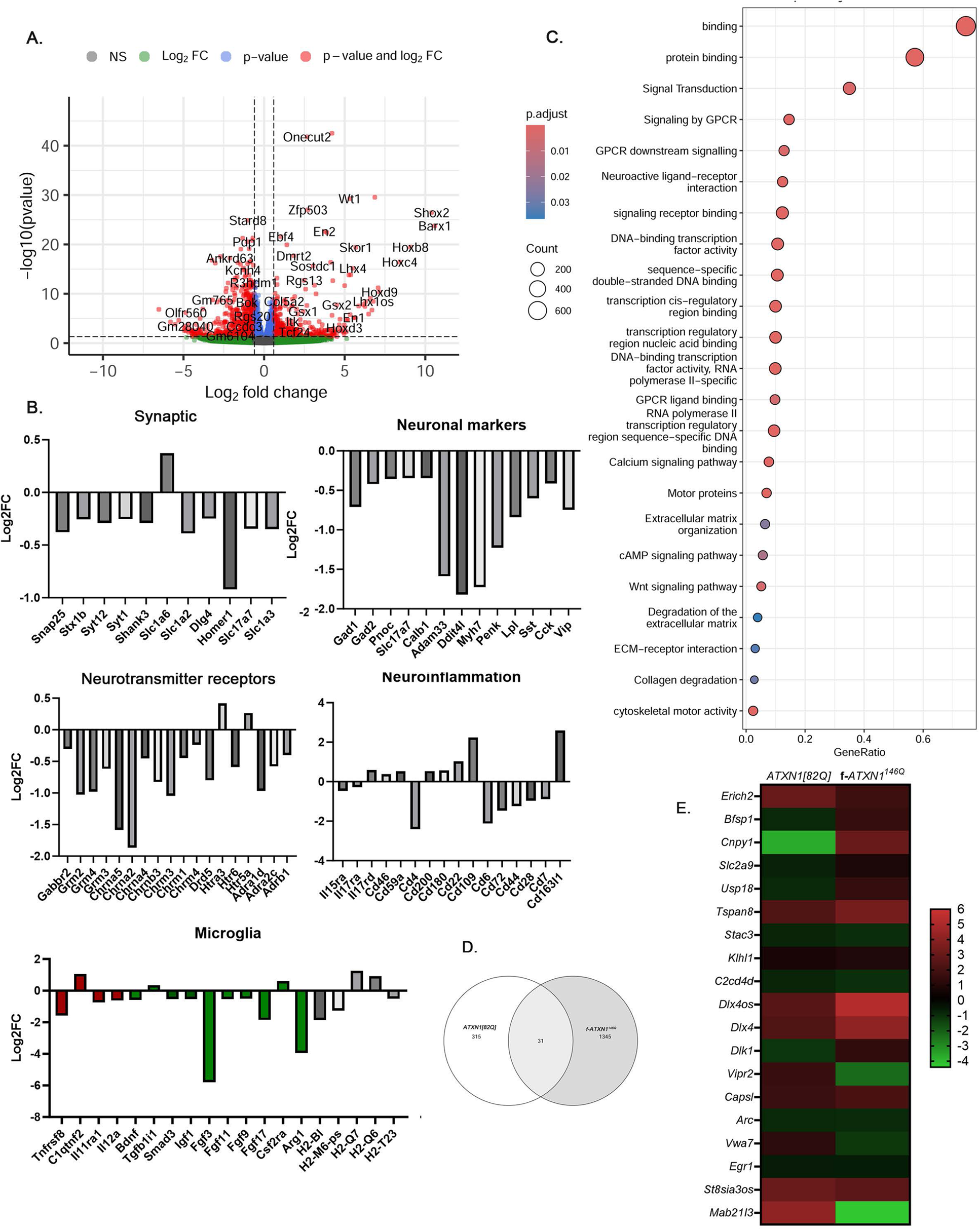
Altered gene expression in *f-ATXN1^146Q^* mice. A. Volcano Plot of PFC Differentially Expressed genes (DEGs) between WT and *f-ATXN1^146Q^* mice ( N= 6 each). B. Expression of synaptic, markers of different neuronal cell types, neurotransmitter receptors, neuroinflammatory and microglial genes is altered in the PFC of *f-ATXN1^146Q^* mice. For all genes p < 0.05. C. Pathway analysis indicating altered signaling in PFC of *f-ATXN1^146Q^* mice. D. Overlap of PFC significant DEGs between WT and *ATXN1[82Q]* mice and WT and *f-ATXN1^146Q^* mice. E. Heatmap of DEGs DEGs between WT and *ATXN1[82Q]* mice and WT and *f-ATXN1^146Q^* mice.

GO analysis of the upregulated DEGs identified altered DNA-binding transcription factor activity and regulation of transcription by RNA polymerase II. Reactome analysis (REAC) identified ECM pathways including assembly of collagen and non-integrin membrane-ECM interaction. GO, KEGG and REAC analysis of the downregulated DEGs identified many signaling pathways including signaling receptor binding, calcium ion binding, ECM structural constituent, neuropeptide signaling pathway, neuroactive ligand-receptor interaction, Wnt, GPCR, and calcium signaling pathways (Figure 4C).

### Arc is altered in PFC of both PC-transgenic and knock-in SCA1 mice

Comparing transcriptional changes in the PFC of *ATXN1[82Q]* and *f-ATXN1^146Q^* mice, we found 31 shared DEGs (Table 3), with approximately half (16 DEGs) changed in the same direction (Figure 4D). Among these congruent genes was immediate-early gene *Arc* that was decreased in the PFC of both *ATXN1[82Q]* and *f-ATXN1^146Q^* mice (Figure 4E). *Arc* is essential for long-lasting information storage in the mammalian brain, mediates various forms of synaptic plasticity, and has been implicated in neurodevelopmental disorders and cognitive deficits in Alzheimer’s disease^47–52^. Another immediate early gene similarly altered in the PFC of *ATXN1[82Q]* and *f-ATXN1^146Q^* was another *Erg1*, also implicated in synaptic function, immune response and cognitive dysfunction^53^.

These results characterize PFC synaptic and gene expression dysfunction in SCA1 mice, and identify several genes altered in PFC of both *ATXN1[82Q]* and *f-ATXN1^146Q^* mice.

### Deleting expanded ATXN1 in PCs exacerbates PFC dysfunction and cognitive impairments in SCA1

It is likely that cerebellar PC dysfunction contributes to altered expression of genes such as Arc and Erg1 that are impacted congruently in *ATXN1[82Q]* and *f-ATXN1^146Q^* mice. To direct test the role of expanded ATXN1 expression in PCs on transcriptional alterations in SCA1, we depleted expanded ATXN1 expression only in PCs of SCA1 mice. In *f-ATXN1^146Q^* mice, mutant human *ATXN1^146Q^* has LoxN recombination sites flanking the coding exons such that the human coding exons of the expanded *ATXN1^146Q^* can be deleted in the presence of Cre recombinase^27^. To accomplish this, we crossed conditional *f-ATXN 1^146Q/2Q^* mice with a *Pcp2-Cre* line that expresses Cre-recombinase only in PCs^28^, and performed RNAseq on the PFC dissected from *f-ATXN1^146Q^;Pcp2-Cre* mice. We identified 1241 significant DEGs in PFC from *f-ATXN1^146Q^;Pcp2-Cre* mice compared to their WT littermates, with similar numbers of upregulated (598) and downregulated (643) DEGs (Figure 5A, Table 4). When comparing DEGs between WT and *f-ATXN1^146Q^* and DEGs between WT and *f-ATXN1^146Q^;Pcp2-Cre* mice, we found 749 shared DEGs and all were altered in the same direction (Table 5, Figure 5B). These included genes encoding for synaptic proteins, neurotransmitter receptors, neuropeptide binding and calcium signaling (Figure 5C). GO, KEGG and REAC pathway analysis of these shared DEGs identified protein binding, signaling by GPCR, calcium signaling, calcium ion binding, and synapses. Noted differences in affected pathways between *f-ATXN1^146Q^* and *f-ATXN1^146Q^;Pcp2-Cre* mice were Wnt and ECM pathways that were altered only in *f-ATXN1^146Q^* mice and glutamatergic synapses pathway that was uniquely altered in *f-ATXN1^146Q^;Pcp2-Cre* mice (Figure 5D). Additionally, calcium signaling and signaling by GPCR were more significantly impacted in *f-ATXN1^146Q^;Pcp2-Cre* mice, while DNA binding transcription factor activity and cytoskeletal motor activity were more significantly altered in *f-ATXN1^146Q^* mice (Supplementary Figure 5).

**Figure 5.**
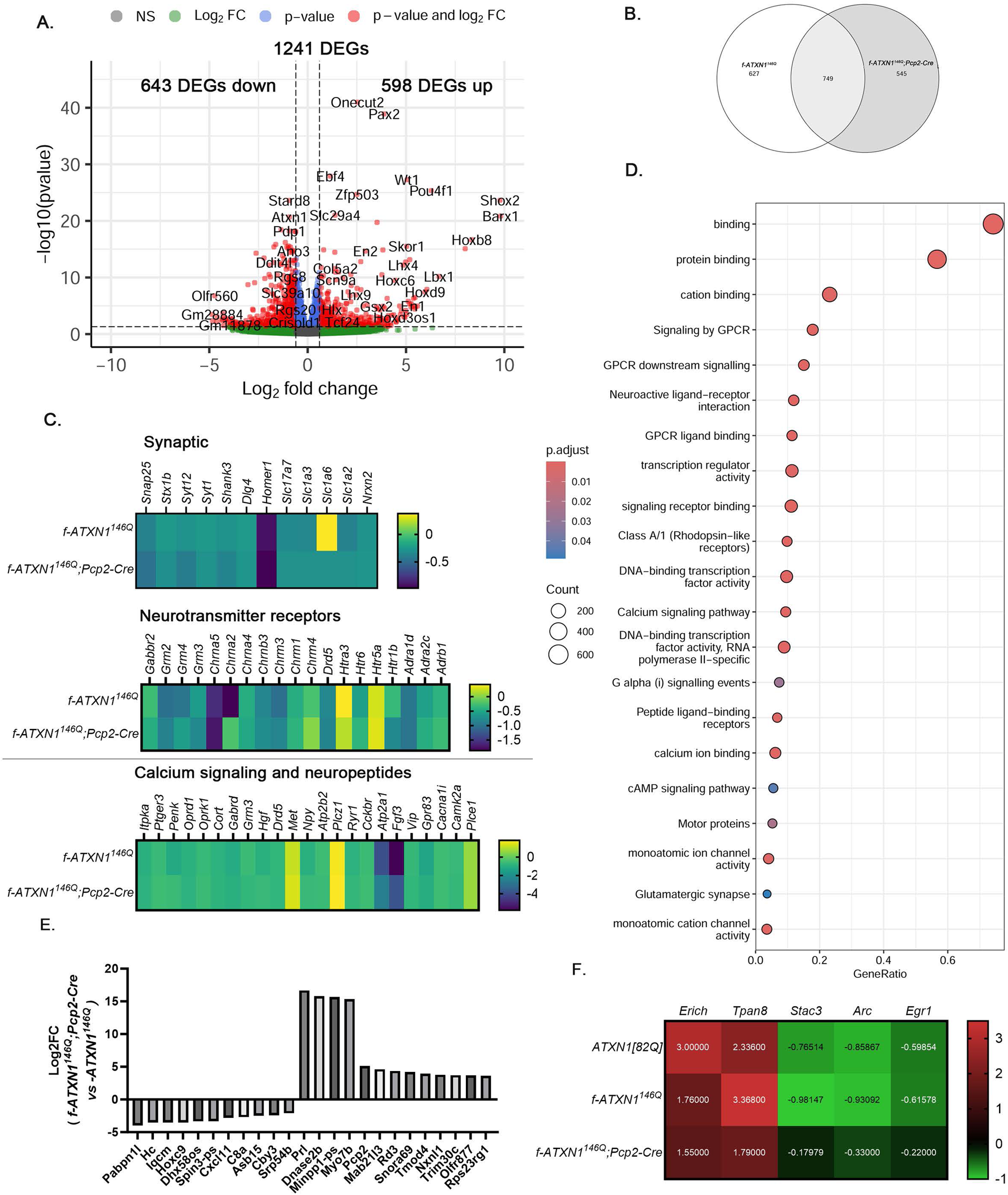
Deleting expanded ATXN1 in PCs impacts gene expression in SCA1 PFC. A. Volcano Plot of PFC Differentially Expressed genes (DEGs) between WT and *f-ATXN1*^146^*^Q^;Pcp2-Cre* mice ( N= 6 each). B. Overlap of PFC significant DEGs between WT and *f-ATXN1^146Q^* and between WT and *f-ATXN1*^146^*^Q^; Pcp2-Cre* mice. C. Shared DEGs between WT and *f-ATXN1^146Q^* and between WT and *f-ATXN1*^146^*^Q^; Pcp2-Cre* mice. For all genes p < 0.05. D. Pathway analysis indicating altered signaling in PFC of *f-ATXN1*^146^*^Q^;Pcp2-Cre* mice. E. Top 10 upregulated and downregulated DEGs between of *f-ATXN1^146Q^* and *f-ATXN1*^146^*^Q^;Pcp2-Cre.* F. Heatmap of PFC DEGs between WT and *ATXN1[82Q]* mice and WT and *f-ATXN1^146Q^* mice, whose expression is normalized in *f-ATXN1*^146^*^Q^; Pcp2-Cre* PFC.

When comparing gene expression between f*-ATXN1^146Q^* and *f-ATXN1^146Q^;Pcp2-Cre* mice we identified 393 DEGs. The top ten upregulated and downregulated DEGs in this comparison (Figure 5E, Table 6) include *Pabnl1* encoding for poly(A) binding protein involved in mRNA decay^54^*, Hc* and *C8a* encoding for complement proteins, *Cxcl11* encoding for chemokine, *Prl* encoding for prolactin that plays a role in synaptic transmission and regulating microglial inflammation^55^, and *Dnase2* involved in interferon signaling^56^. Intriguingly, 208 DEGs in this comparison are not significantly altered in either WT vs f*-ATXN1^146Q^* or WT vs *f-ATXN1^146Q^;Pcp2-Cre* comparisons. Expression of 120 DEGs (∼ 30%) are significantly altered in *f-ATXN1^146Q^* vs WT comparison, and are either not significantly altered (99 DEGs) or the change is ameliorated/normalized (21 DEGs) in *f-ATXN1^146Q^;Pcp2-Cre* mice. Transcription factor analysis of 90 DEGs that were downregulated in *f-ATXN1^146Q^* mice, but not altered in *f-ATXN1^146Q^;Pcp2-Cre* mice identified *Foxn2* as a transcriptional factor. 26 and 27 DEGs were upregulated or downregulated only in *f-ATXN1^146Q^;Pcp2-Cre* compared to their WT littermates (i.e. not altered in WT vs *f-ATXN1^146Q^* PFC). Among genes upregulated only in *f-ATXN1^146Q^;Pcp2-Cre* mice were *Fas, Itgax, Csf2rb, CCL3, IL18rap,* encoding for inflammatory and cell-death proteins. Among genes downregulated only in *f-ATXN1^146Q^;Pcp2-Cre* mice were *Pabnl1*,*CD209g* and *Kbtbd6* involved in mRNA decay, carbohydrate binding and ubiquitination.

We next wanted to examine which of the genes that are altered in the same direction in *ATXN1[82Q]* and *f-ATXN1^146Q^* PFC had their expression normalized in f*-ATXN1^146Q^;Pcp2-Cre* mice. We found that expression of *Arc, Egl1*, *Erich, Tpan8 a*nd *Stac3* were normalized in *f-ATXN1^146Q^;Pcp2-Cre* mice, indicating that expanded ATXN1 in PCs may contribute to their altered expression in SCA1 knock-in and transgenic mice (Figure 5F). Together, these results indicate that expression of expanded ATXN1 in Purkinje cells impacts gene expression changes in the PFC of SCA1 mice. Expression of some genes was normalized by deletion of expanded ATXN1 in Purkinje cells. Depending whether these genes play cause or compensate for PFC pathology and behavioral deficits, normalizing their expression may ameliorate or exacerbate behavioral deficits.

### Deleting expanded ATXN1 in PCs exacerbates performance of SCA1 mice on Barnes Maze and decreases their enhanced freezing in fear conditioning

To determine the effect of expanded ATXN1 expression in PCs on SCA1-like cognitive deficits we compared the performance of *f-ATXN1^146Q^*, f*-ATXN1^146Q^;Pcp2-Cre,* and WT littermates on Barnes Maze and Fear conditioning (FC). Mice were first tested on Barnes Maze and FC at 12 and 13 weeks respectively, then we tested their memory recall at 25 and 26 weeks, which was followed by retraining (Figure 6A).

**Figure 6.**
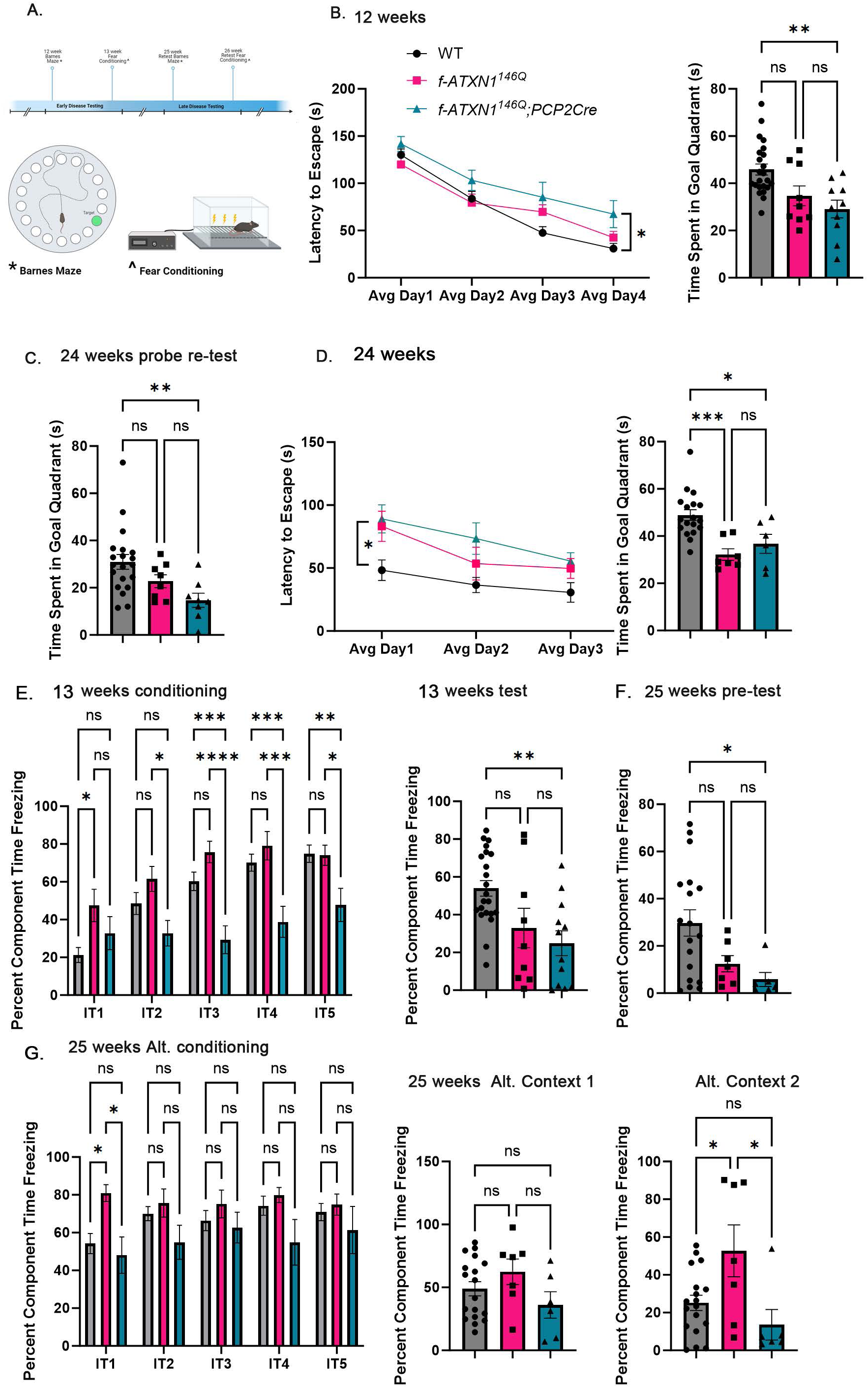
Altered cognitive performance on Barnes Maze and Fear Conditioning of *f-ATXN1^146Q^; Pcp2-Cre* mice. A. Schematic illustrating behavioral testing. B. Barnes Maze training and testing at 12 weeks. C. Retesting at 24 weeks. D. Barnes Maze training and testing at 24 weeks. E. Fear conditioning and testing next day at 13 weeks. F. Fear conditioning pre-testing at 25 weeks. G. Fear conditioning with alternative context 1 and testing next day at 25 weeks. N= 24, 9, and 10 WT, *f-ATXN1^146Q^* and in *f-ATXN1*^146^*^Q^; Pcp2-Cre* mice. Data is mean ± SEM with dots representing individual mice. Two-Way ANOVAs was used to analyze training on Barnes Maze and conditioning on Fear Conditioning. One-Way ANOVA was used for testing and re-testing. P values represent post-hoc Tukey test. * p < 0.05. ** p< 0.01, *** p< 0.005, **** p< 0.0001.

For Barnes Maze, mice underwent four days of training to learn how to find the escape hole at 12 weeks. With each day of training, all groups of mice showed a decrease in the average latencies to find the hole, indicative of learning (Figure 6B). However, while latencies during training were indistinguishable for *f-ATXN1^146Q^* and WT mice, latencies for *f-ATXN1^146Q^;Pcp2-Cre* mice were significantly longer. During the probe day, when there was no escape hole, time spent in the goal quadrant was not statistically different for WT and *f-ATXN1^146Q^*. Time in the goal zone was significantly reduced for *f-ATXN1^146Q^;Pcp2-Cre* mice compared to WT mice (Figure 6B). We used Barnes maze unbiased strategy (BUNS) analysis^26,57^ to determine which strategies are mice using to find the escape hole. With each day of training, WT mice increasingly used spatial strategies to find the escape hole. However, both *f-ATXN1^146Q^* and *f-ATXN1^146Q^;Pcp2-Cre* mice were preferentially using serial strategy ( Supplementary Figure 6), and as a result both had lower cognitive score. These results indicate that early in disease, while *f-ATXN1^146Q^* mice are undistinguishable from WT mice on latencies and time in goal quadrant during learning and memory testing respectively, they do show deficit in strategy development. Moreover, depleting expanded ATXN1 from PCs exacerbated both learning and memory of *f-ATXN1^146Q^;Pcp2-Cre* mice on the Bares Maze at this early disease stage.

To determine how disease progression and expanded ATXN1 in PCs impact memory recall and cognitive performance of *f-ATXN1^146Q^* mice we repeated Barnes Maze at 25 weeks. We first performed a pre-test to evaluate memory recall (i.e. whether mice can remember where the escape hole was during training at 12 weeks). While *f-ATXN1^146Q^* and WT mice were indistinguishable in the amount of time in the goal quadrant, *f-ATXN1^146Q^;Pcp2-Cre* mice had a significantly lower time in the goal quadrant (Figure 6C). To test whether SCA1 mice have impaired ability to learn at this later disease stage mice were re-trained on Barnes Maze for three days (Figure 6D). At 24 weeks, both *f-ATXN1^146Q^* and for *f-ATXN1^146Q^;Pcp2-Cre* showed increased latencies during training compared to WT mice, and both were impaired on the probe test.

We also performed a fear conditioning assay at 13 weeks. During the first bout of conditioning (IT1), *f-ATXN1^146Q^* were freezing significantly more than WT mice and percentage freezing time for *f-ATXN1^146Q^;Pcp2-Cre* was in-between. Both WT and *f-ATXN1^146Q^* mice progressively increased freezing in response to repeated shock (IT1-IT5) demonstrating learning of fear response. Surprisingly, this was absent in *f-ATXN1^146Q^;Pcp2-Cre* mice and they were freezing significantly less compared to WT and *f-ATXN1^146Q^* at each of subsequent trials (IT2-IT5) (Figure 6E). The next day, mice were tested for freezing response in the absence of shock. *f-ATXN1*^146^*^Q^;Pcp2-Cre* mice were still freezing significantly less compared to WT mice (Figure 6F).

Mice underwent a second FC at 26 weeks starting with a pre-test to evaluate recall. While both *f-ATXN1^146Q^* and *f-ATXN1*^146^*^Q^;Pcp2-Cre* were freezing less compared to WT mice, only *f-ATXN1*^146^*^Q^;Pcp2-Cre* were significantly different (Figure 6G). During FC in the alternative context 1 at 26 weeks, *f-ATXN1^146Q^* were freezing more during IT1 but at IT2-IT5 all three groups of mice were indistinguishable. On the probe test the next day, we did not find a significant difference in freezing among three groups in the alternative context 1. Surprisingly, *f-ATXN1^146Q^* were freezing more in the alternative context 2 that was not used for FC. These results indicate an enhanced freezing response in *f-ATXN1^146Q^* mice compared to WT mice. Deleting expanded ATXN1 from PCs resulted in a lack of freezing response in SCA1 mice, suggesting that cerebellar dysfunction contributes to the freezing response in SCA1.

Together, these results indicate that cerebellar dysfunction contributes to cognitive and mood deficits in SCA1 mice. Removal of expanded ATXN1 from PCs exacerbated cognitive decline seen in SCA1 mice, implying that cerebellar dysfunction may compensate for PFC pathology and revealing a more complex relationship between the brain regions in disease.

## Discussion

One of the key questions in neurodegeneration is understanding the similarities and differences in how different brain regions and cell types are impacted in disease. Building on top of this question is increasing our understanding of how different brain regions affected in disease impact on each other to produce disease symptoms.

Our previous study demonstrated that expressing expanded ATXN1 only in cerebellar PCs is sufficient to cause cognitive and mood deficits in PC transgenic *ATXN1[82Q]* mice^26,58^. This is consistent with cerebellar cognitive affective syndrome (CCAS) also known as Schmahmann’s syndrome and in patients with cerebellum limited strokes and degeneration. These patients demonstrate impaired cognition, particularly in executive function related to the PFC^6,59^. Cerebello-cerebral anatomical and functional connectivity has been proposed as the mechanism by which the cerebellum impacts cognition and affect^6,7,9^. What was unclear is how cerebellar dysfunction impacts the PFC at cellular and molecular level. We demonstrated here reduced c-Fos activity, decrease in synapses, and gene expression changes in *ATXN1[82Q]* mice. These results indicatethat cerebellar dysfunction via cerebello-cerebral loops is sufficient to alter expression of key immediate early genes in the PFC, reduce neuronal activity and synaptic density.

We have also found that cognitive deficits are different in *Atxn1^154Q/2Q^* mice expressing mutant ATXN1 throughout the brain. This suggested more complex cerebello-cerebral pathology in mice globally expressing expanded ATXN1 that impacts their cognitive performance differently than in PC specific *ATXN1[82Q]* mice^26,58^. Indeed, early in disease we found increased neuronal and synaptic density in the PFC of *f-ATXN1^146Q^* mice, a novel conditional SCA1 knock-in model^27^. At a later disease stage, we found a decrease in c-Fos and synaptic density indicative of progressive PFC pathology. We found a large number of DEGs, indicating significant synaptic, calcium, and neuroinflammatory alterations in the PFC of *f-ATXN1^146Q^* mice. Importantly, we have found a decrease in the expression of genes encoding for cholinergic receptors, consistent with decrease seen in patients with SCA1^60^. We found 16 genes that were altered in the PFC of both *ATXN1[82Q]* and *f-ATXN1^146Q^* mice. Importantly, immediate early genes Arc and Erg1 are decreased in the PFC of both mice. As both Arc and Erg1 are known to play key roles in synapses, cognition, and mood, we speculate that this decrease in their expression may contribute to PFC dysfunction and impaired cognition and mood in SCA1 transgenic and knock-in mice.

To determine how expanded ATXN1 expression in PCs impacts cognitive performance and PFC dysfunction we created *f-ATXN1^146Q^*;*Pcp2-Cre* in which mutant ATXN1 is deleted only in PCs using a cre-lox system. Surprisingly, PFC pathology and cognitive performance on Barnes Maze were exacerbated in *f-ATXN1^146Q^*;*Pcp2-Cre* mice, indicating that PC dysfunction may have a beneficial impact on cognition during early stages of disease. On the other hand, the exaggerated freezing response seen in *f-ATXN1^146Q^* mice was fully reversed in *f-ATXN1^146Q^*;*Pcp2-Cre* mice who did not demonstrate significant freezing early in disease. This may indicate that PC dysfunction contributes to increased freezing or anxiety seen throughout disease in SCA1 mice. Expression of both *Arc* and *Erg1* was ameliorated in the PFC of *f-ATXN1^146Q^*;*Pcp2-Cre*, further supporting that PC dysfunction contributes to decreased *Arc* and *Erg1* expression and increased freezing of *f-ATXN1^146Q^* mice. Future studies could rescue Arc and/or Erg1 expression in the PFC to determine their role in SCA1 mood alterations.

We have also found 121 DEGs whose expression is altered in *f-ATXN1^146Q^* mice and normalized in *f-ATXN1^146Q^*;*Pcp2-Cre* mice, indicating that expanded ATXN1 in PCs contributes to their altered expression. On the other hand we found 53 DEGs whose expression is altered only in *f-ATXN1^146Q^*;*Pcp2-Cre* mice but not in *f-ATXN1^146Q^* mice. These results further support a complex cerebello-cerebral interplay in SCA1 pathogenesis, revealing that pathology in one region may have a beneficial effect on another region.

There are several important implications of our findings. First, circumscribed dysfunction in one brain region, such as caused by expression of expanded ATXN1 only in Purkinje cells in *ATXN1[82Q]* mice is sufficient to alter synaptic properties and gene expression in another brain region, such as prefrontal cortex. Second, when both prefrontal cortex and cerebellum are undergoing pathology, cerebellum may may have a beneficial influence on prefrontal cortex, implicating that the impact of cerebellar dysfynction on prefrontal cortex is context dependent. It is possible that cerebellum has similar effect on other cortical or subcortical regions that it communicates with and will be examined in the future. Finally, we demonstrated differing effects of deleting expanded ATXN1 in PCs on different aspects of cognition. Deleting expanded ATXN1 in PCs exacerbated learning and memory on Barnes Maze, implicating that PC-expression of expanded ATXN1 ameliorates spatial learning and memory aspects of SCA1-like cognitive deficits. On the other hand, deleting expanded ATXN1 in PCs reversed mouse behavior on Fear Conditioning from exacerbated freezing seen in *f-ATXN1^146Q^* mice compared to WT controls, to almost no freezing in *f-ATXN1^146Q^*;*Pcp2-Cre* mice, implicating that PC-expression of expanded ATXN1 contributes to increased anxiety/freezing response aspects of SCA1-like cognitive deficits. Future studies will determine the role of altered Arc expression in enhanced freezing response in SCA1. We hope that other DEGs and pathways that we identified will inspire additional work on understanding cognitive deficits and regional communication in disease.

## Data availability

All the data and materials will be available upon contacting the authors for requests.

## Acknowledgements

We acknowledge all the members of the Orr and Cvetanovic laboratories for thoughtful discussions and feedback on the study. Work in this study was aided by the Mouse Behavioral Core and Genomics Core at the University of Minnesota. This work was supported by National Institute of Health NINDS award (R01 NS197387 to M.C.).

---

**Supplementary Figure 1. Illustrating the location of PFC immunohistochemistry.**

**Supplementary Figure 2. Activity of PFC neurons is altered in *ATXN1[82Q]* mice at 18 weeks.** Slices from 18 weeks old ATXN1[82Q] mice and wild-type littermate controls were stained using Ab against markers of neurons (NeuN), activity (c-Fos), excitatory synapses(VGLUT2 and PSD95), and inhibitory synapses (VGAT and gephyrin). Confocal z-stacks were used to quantify neuronal density (A), activity (B), pre-and postsynaptic VGLUT2 and PSD 95 puncta and their co-localization (C) and pre and postsynaptic VGAT and Gelhyrin puncta and their colocalization (D). Data is mean ± SEM with dots presenting individual mice or brain sections. N = 6-12 mice of each genotype. Student’s test *** p< 0.0005.

**Supplementary Figure 3. *Atxn1* expression in different brain regions.** We plotted *Atxn1* expression as Log of Counts per Million (CPM) in cortex, striatum, hippocampus, cerebellum and brain stem using our RNAseq data^45^

**Supplementary Figure 4. Figure 3. PFC pathology in *f-ATXN1^146Q^* mice.** Saggital brain sections of 28-week-old *f-ATXN1^146Q^* mice and their wild-type littermates were immunostained for NeuN, c-Fos, VGLUT2, PSD95, VGAT and gephyrin, and confocal z-stack images were obtained. A. Quantification of NeuN density. B. Quantification of c-Fos density. C. Image of VGLUT2 and PSD95 puncta. D. Quantification of excitatory presynaptic VGLUT2+, postsynaptic PSD95+ and co-localized VGLUT2/PSD95 puncta. E. Quantification of inhibitory presynaptic VGAT+, postsynaptic Gephyrin+ and co-localized VGAT/gephyrin puncta. N= 6-12 *f-ATXN1^146Q^* mice and their wild-type littermates. * P< 0.05, ** P<0.01 Student’s t-test.

**Supplementary Figure 5. KEGG pathways analysis of DEGs in WT vs ATXN1[82Q], WT vs *f-ATXN1***^146^***^Q^* and WT vs. *f-ATXN1***^146^***^Q^; Pcp2-Cre* PFC comparisons.**

**Supplementary Figure 6. BUNS analysis of strategy development on Barnes maze at 12 and 25 weeks.** Two-way ANOVA was used to analyze cognitive scores.* p<0.05, ** P< 0.01.

