## Supplemental Figures for "Cerebellar contribution to cognitive deficits and prefrontal cortex dysfunction in Spinocerebellar Ataxia Type 1 (SCA1)"

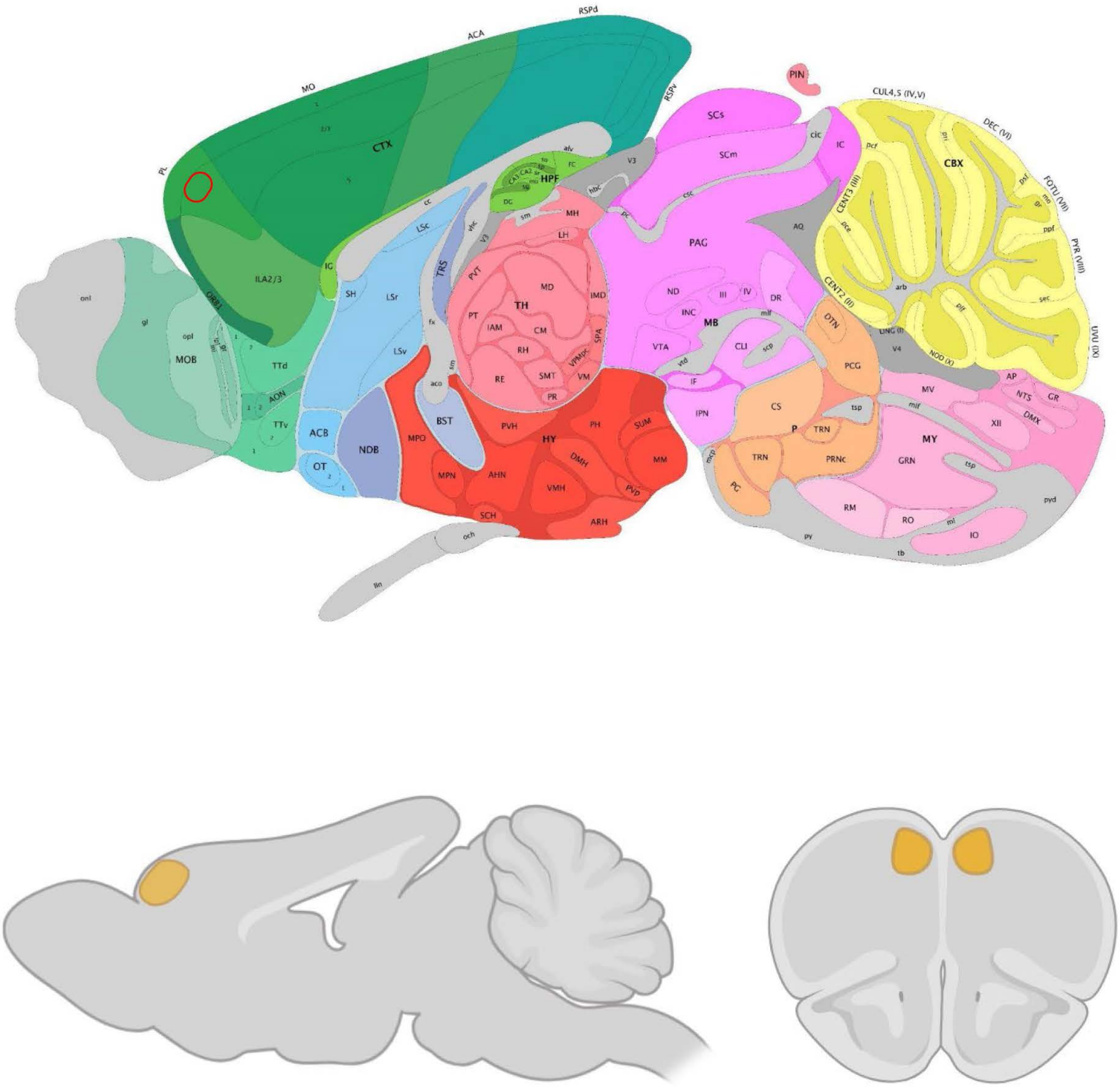


Supplementary Figure 1.

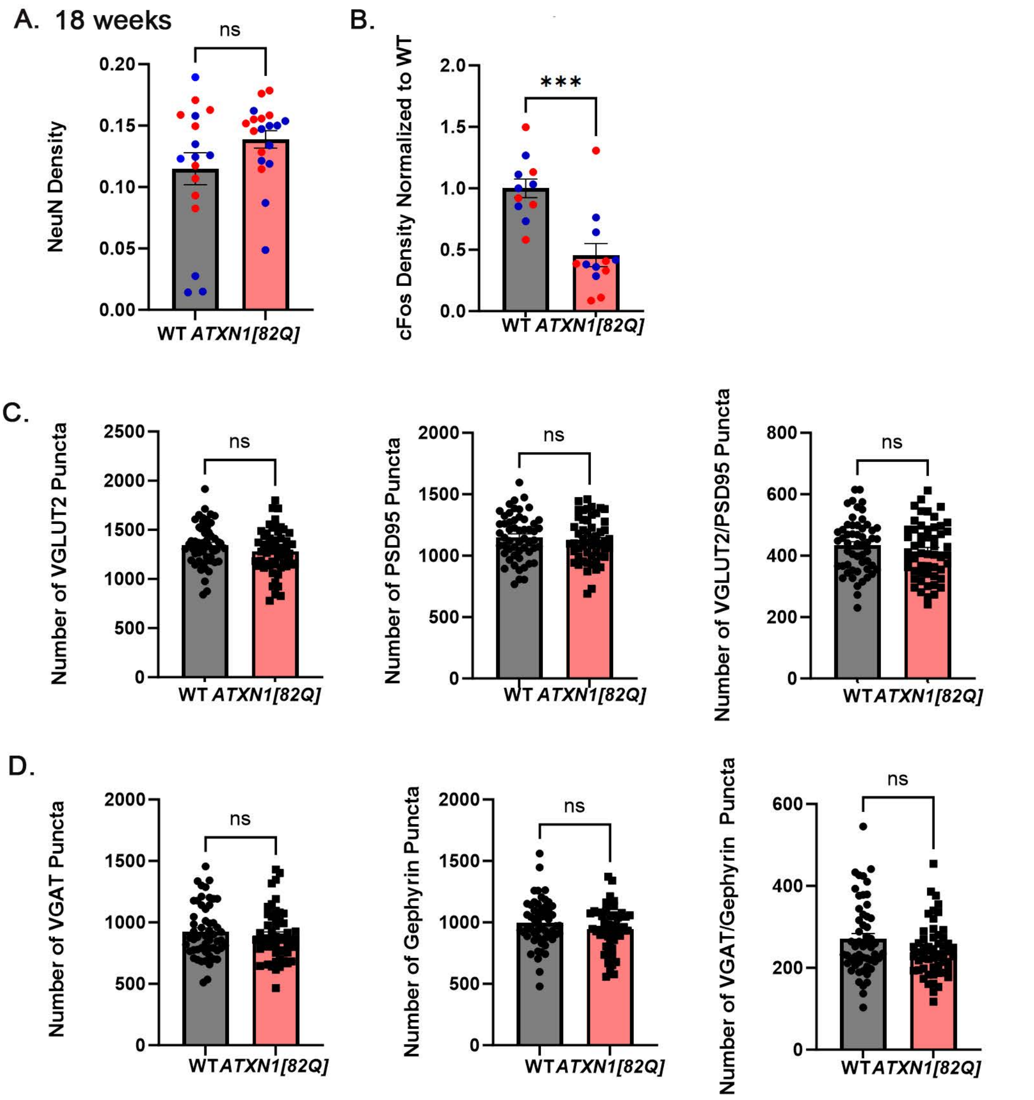


Supplemetary Figure 2.

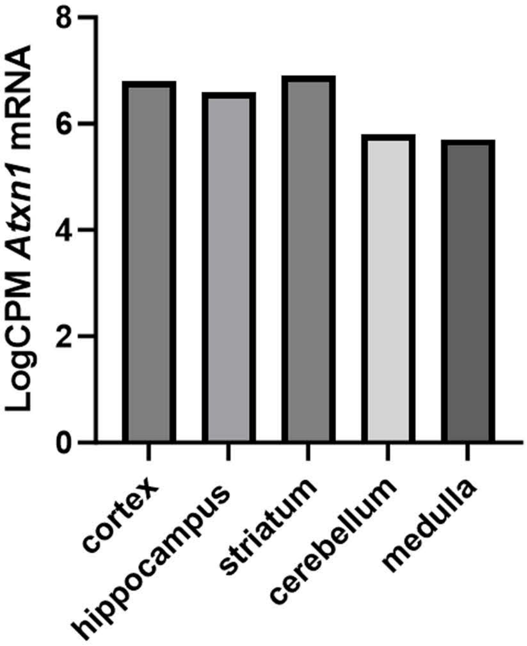


Supplementary Figure 3.

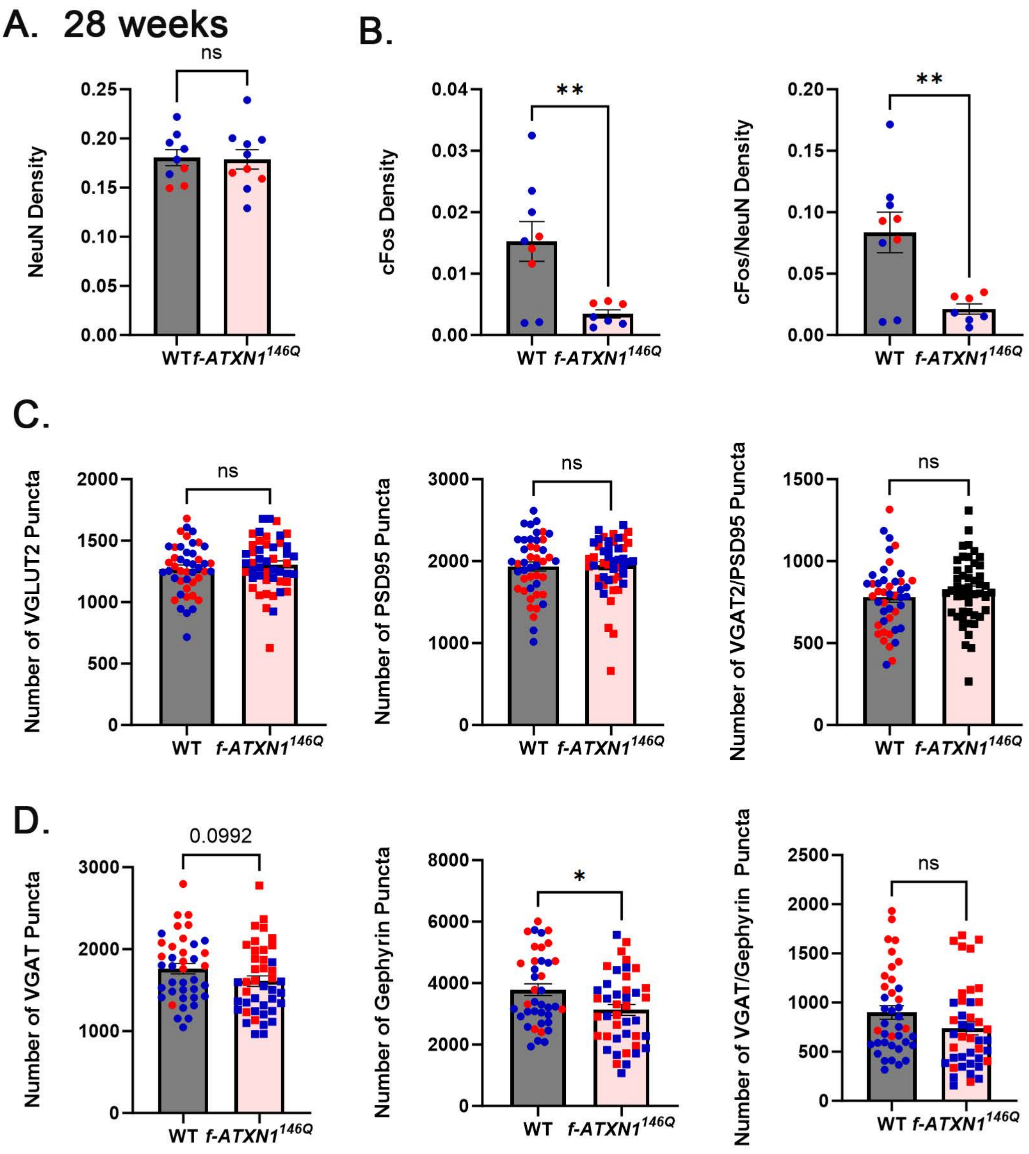


Supplementary Figure 4.

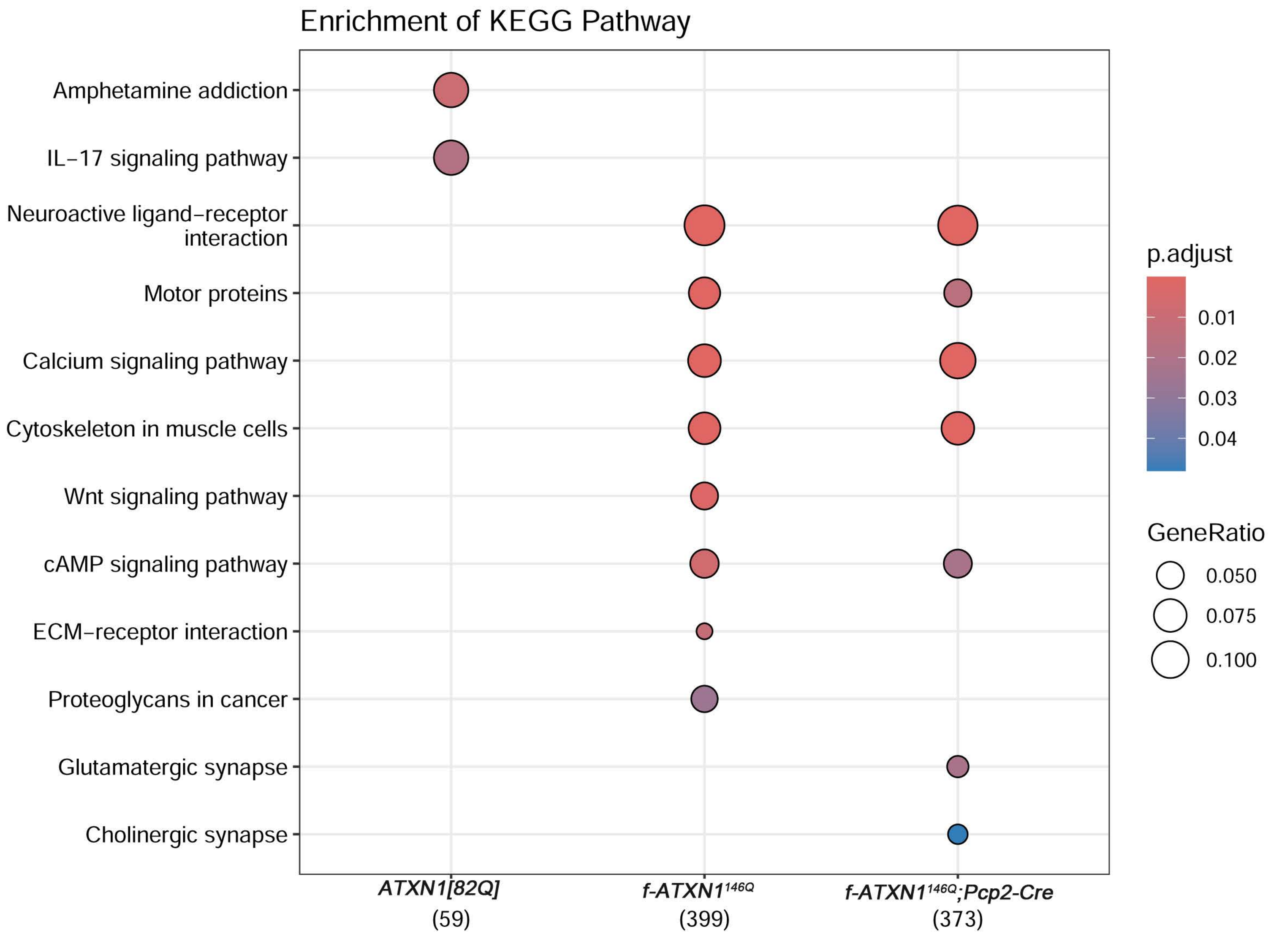


**Supplementary Figure 5.**

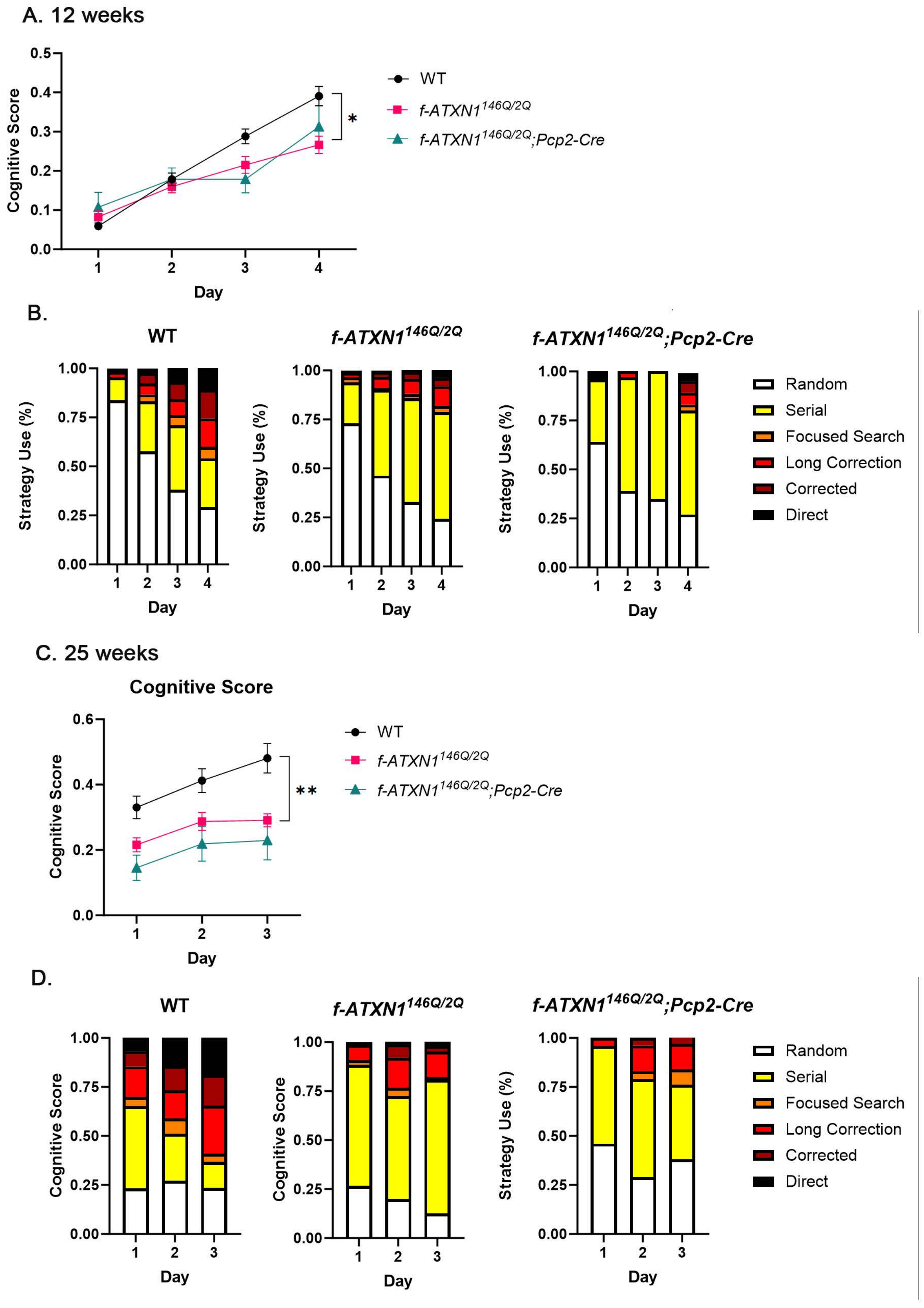


Supplementary Figure 6.
